# Absence of SIV-specific CD8+ T cells in germinal centers may set the stage for persistent infection

**DOI:** 10.1101/407528

**Authors:** Shengbin Li, Joy M. Folkvord, Katalin J Kovacs, Reece K. Wagstaff, Gwantwa Mwakalundwa, Aaron K. Rendahl, Eva G. Rakasz, Elizabeth Connick, Pamela J. Skinner

## Abstract

CD8+ T cells play an important role in controlling of HIV and SIV infections. However, these cells are largely excluded from B cell follicles where HIV and SIV producing cells concentrate during chronic infection. It is not known, however, if antigen-specific CD8+ T cells are excluded gradually as pathogenesis progresses from early to chronic phase, or this phenomenon occurs from the beginning infection. In this study we determined that SIV-specific CD8+ T cells were largely excluded from follicles during early infection, we also found that within follicles, they were entirely absent in 60% of the germinal centers (GCs) examined. Furthermore, levels of SIV-specific CD8+ T cells in follicular but not extrafollicular areas significantly correlated inversely with levels of viral RNA+ cells. In addition, subsets of follicular SIV-specific CD8+ T cells were activated and proliferating and expressed the cytolytic protein perforin. These studies suggest that a paucity of SIV-specific CD8+ T cells in follicles and complete absence within GCs during early infection may set the stage for the establishment of persistent chronic infection.

**Author Summary:** A paucity of SIV-specific CD8+ T cells in lymphoid follicles and complete absence within most follicular germinal centers during early infection may set the stage for the establishment of persistent chronic infection.

## Introduction

Most human immunodeficiency virus (HIV)-infected individuals fail to adequately control persistent high-level viral replication that results in gradual loss of CD4 T cells and ultimately AIDS in the absence of antiretroviral therapy (ART). B cell follicles in secondary lymphoid tissues have been identified as important sanctuaries that contain large amounts of virus-producing cells during chronic HIV and simian immunodeficiency virus (SIV) infection (Folkvord et al., 2005; Connick et al., 2007, 2014; Hufert et al., 1997; Brenchley et al., 2012). CD4^+^ T follicular helper (T_FH_) cells, a population that mainly resides in B cell follicles, serve as a major site of productive HIV and SIV replication during the chronic phase of infection (Folkvord et al., 2005; Connick et al., 2007; Hufert et al., 1997; Embretson et al., 1993; Perreau et al., 2013; Gratton et al., 2000). In SIV-infected rhesus macaques that control viral replication, either via a natural highly effective immune response or receiving long-term, fully suppressive ART, residual productive SIV infection is strikingly restricted to T_FH_ cells (Fukazawa et al., 2015). In HIV infected aviremic individuals treated with long-term ART, T_FH_ also serves as a major reservoir for active and persistent virus transcription (Banga et al., 2016). Therefore, understanding the immune activity needed to kill virus-infected T_FH_ cells in B cell follicle is necessary for developing novel therapies to fully eradicate HIV or SIV infection.

Antigen-specific CD8+ T cells have a key role in controlling HIV and SIV infections. Their emergence during the acute phase of infection is associated with a decline in plasma viremia (Borrow et al., 1994; Koup et al., 1994; Kuroda et al., 1999). Moreover, the transient depletion of CD8+ T cells during SIV or SHIV infections induces high levels of plasma viremia which are reduced upon reconstitution of CD8+ lymphocytes (Schmitz et al., 1999; Jin et al., 1999; Matano et al., 1998). Strong HIV-specific CD8+ T cell activity is directly associated with long-term elite control of infection (Migueles et al., 2002; Betts et al., 2006; Hersperger et al., 2010). Furthermore, we previously showed a significant inverse relationship between SIV-specific CD8+ T cell frequency and SIV-producing cell levels in lymphoid compartments during chronic SIV infection (Connick et al., 2014). However, in spite of the notable anti-viral effect, HIV- and SIV-specific CD8+ T cells fail to fully eliminate viral replication and the vast majority of HIV and SIV-infected individuals eventually develop disease in the absence of ART.

We and others previously showed that HIV- and SIV-specific CD8+ T cells are largely excluded from B cell follicles in lymph node and spleen tissues during chronic infection (Connick et al., 2007, 2014; Li et al., 2016; Tjernlund et al., 2010). The paucity of virus-specific CD8+ T cells inside B cell follicles, where HIV- and SIV-producing cells are highly concentrated, creates an immune privileged site and an important mechanism of immune evasion by HIV and SIV. This mechanism may, at least partially, account for the failure of CD8+ T cells to fully eradicate HIV/SIV infection.

The exclusion of anti-viral CD8+ T cells from B cell follicles during chronic infection is not absolute. Studies indicate that there are populations of functional CD8+ T cells expressing CXCR5 in B cell follicles in chronic LCMV, HIV and SIV infections (Li et al., 2016; He et al., 2016; Leong et al., 2016), and levels of follicular virus-specific CD8+ T cells correlate with reductions of plasma viral loads and tissue viral replication (Li et al., 2009; Connick et al., 2014; Li et al., 2016; Mylvaganam et al., 2017). Thus, while typically relatively low in numbers, virus-specific CD8+ T cells in follicles appear capable of suppressing viral replication.

Whether virus-specific CD8+ T cells migrate into B cell follicles during early HIV/SIV infection remains to be determined. Our hypothesis is that a paucity of SIV-specific T cells in lymphoid follicles contributes to the establishment of the follicular reservoir of SIV during early SIV infection. To test this hypothesis, in this study, we determined the abundance, distribution and phenotype of SIV-specific T cells in lymph nodes from a cohort of SIV infected rhesus macaques during the early stages of infection.

## Results

### SIV-specific CD8+ T cells are largely excluded from GCs during early infection

To determine whether SIV-specific CD8+ T cells accumulate within B cell follicles during early infection, we evaluated the distribution and quantity of SIV-specific CD8+ T cells at 14 (n=7) and 21 (n=9) days post-infection (dpi) in the lymph nodes from SIV-infected rhesus macaques. All of the animals in this study expressed the MHC-I allele Mamu-A1*001, and immunodominant Gag specific CD8+ T cells were identified using *in situ* tetramer staining with Mamu-A1*001 tetramers loaded with Gag CM9 peptides. In combination with MHC tetramer staining of SIV-specific CD8+ T cells, antibodies against IgM were used to label B cell follicles cells *in situ* in lymph node sections. We detected tetramer+ SIV-specific CD8+ T cells in B cell follicles at both 14 and 21 days post-infection (Fig. 1A). Similar to chronic infection (Connick et al., 2014), the number of SIV-specific tetramer+ CD8+ T cells/mm^2^ inside B cell follicles was significantly lower than in extrafollicular regions at 14 dpi (Fig. 1B) and at 21 dpi (Fig. 1C). However, the extent of this difference was greater during chronic infection which showed a median of 4X lower cells/mm^2^ in follicles (Connick et al., 2014), compared to a median of 2X lower found here at 14 and 21 dpi. We previously showed that there is a positive correlation between numbers of follicular and extrafollicular SIV-specific tetramer+ CD8+ T cells/mm^2^ (Connick et al., 2014; Li et al., 2016). In this study, we investigated the relationship between levels of SIV-specific tetramer+ CD8+ T cells inside and outside B cell follicles during early infection at 21 dpi. A highly significant positive correlation between levels of follicular and extrafollicular SIV-specific tetramer+ CD8+ T cells in lymph node was observed on 21 dpi (Fig. 1D). A similar correlation was seen at 14 dpi (data not shown). These data demonstrated that a small number of SIV-specific CD8+ T cells enter the B cell follicles during early SIV infection and that levels of follicular SIV-specific CD8+ T cells positively correlate to levels of SIV- specific CD8+ T cells in extrafollicular regions.

**FIG 1.**
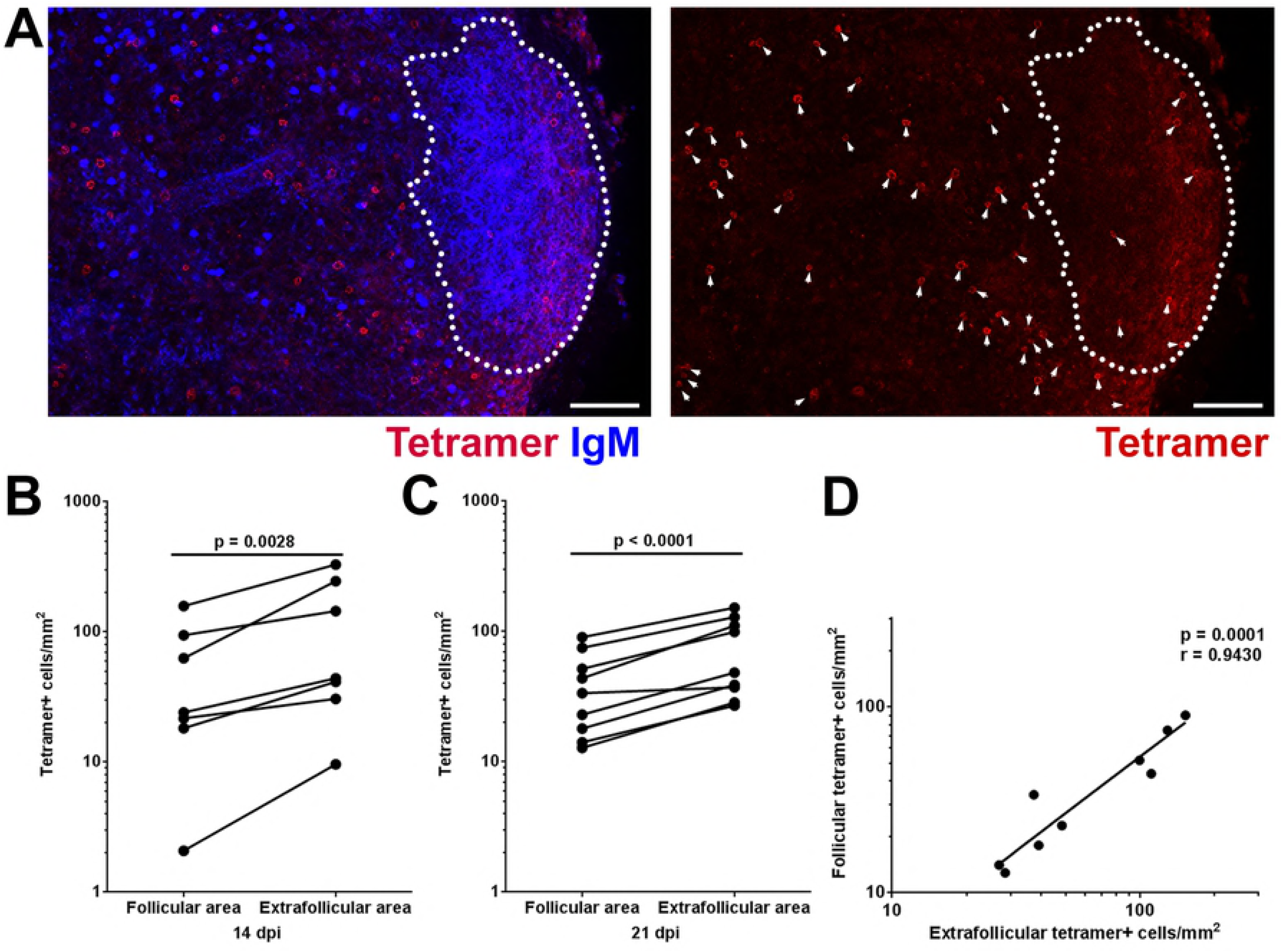
Relatively low levels of SIV-specific CD8 T cells detected in B cell follicles during early SIV infection. (**A**) Representative lymph node section shows the distribution of SIV-specific CD8 T cells in different compartments of lymph node on 21 dpi. This section was stained with Mamu-A1*001/Gag CM9 tetramers to label SIV-specific CD8 T cells (red, and indicted with arrows in the image on the right which shows tetramer staining alone), IgM antibodies (blue) to define follicles (indicated with dotted line). Confocal images were collected with a 20X objective and each scale bar indicates 100 µm. (**B**) Frequencies of tetramer^+^ SIV-specific CD8 T cells in B cell follicle were 57% (95% confidence interval [CI]: 34%, 71%) lower than those in extrafollicular region on 14 dpi (p = 0.0028). (**C**) Frequencies of tetramer^+^ SIV-specific CD8 T cells in B cell follicle were 47% (95% CI: 36%, 56%) lower than those in extrafollicular region (p < 0.0001) on 21 dpi. (**D**) Relationship between frequencies of follicular and extrafollicular tetramer^+^ SIV-specific CD8 T cells. The frequency of follicular tetramer^+^ SIV-specific CD8 T cells is significantly correlated with those located in extrafollicular area (r = 0.943, p = 0.0001); a 1-log increase in extrafollicular region is associated with a 0.99-log increase in follicular region (95% CI: 0.68-log, 1.30-log).

We next analyzed whether SIV-specific tetramer+ CD8+ T cells accumulate within the germinal center (GC) area of the follicles, where follicular dendritic cells (FDCs) retain infectious virus trapped in immune complexes (Heath et al., 1995). To address this question, we used Gag CM9 tetramers to detect SIV-specific CD8+ T cells, IgM-specific antibodies to identify B cell follicles and Ki67-specific antibodies to identify GCs. At 14 dpi, very few lymph node follicles with GCs were detected. At 21 dpi follicles with GC were abundant. Strikingly, at 21 dpi, 60.3% (44/73) of the GCs were completely devoid of SIV-specific tetramer+ CD8+ T cells (Fig. 2A). Furthermore, quantitative analysis showed that the frequency of SIV-specific tetramer+ CD8+ T cells in GCs was significantly lower than non-GC follicular area (Fig. 2B).

**FIG 2.**
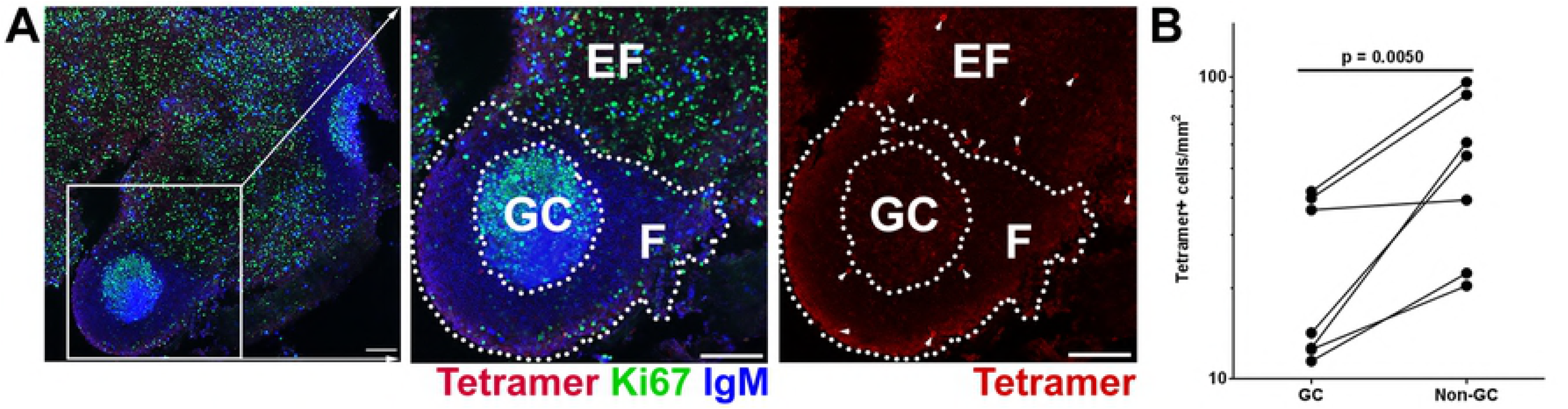
Follicular SIV-specific CD8 T cells were largely excluded from germinal centers (GC) during early SIV infection. (**A**) Representative images demonstrate the distribution of SIV-specific CD8 T cells within B cell follicle during early SIV infection. Sections were stained with Mamu-A1*001/Gag CM9 tetramers to label SIV-specific CD8 T cells (red, and indicted with arrows in the image on the right which shows tetramer staining alone), IgM antibodies (blue) to define follicles (F), and Ki67 antibodies (green) to label GC. Confocal images were collected with a 20X objective and each scale bar indicates 100 µm. (**B**) Frequencies of follicular tetramer^+^ SIV-specific CD8 T cells in GCs were 56% (95% CI: 30%, 73%) lower than those in non-GC follicular area (p = 0.005).

### Many follicular SIV-specific CD8+ T cells express PD-1 in early SIV infection

PD-1 is a marker of functional exhaustion (Barber et al., 2006; Day et al., 2006) as well as a marker of CD8+ T cells that have recently been exposed to antigenic stimulation (Agata et al., 1996). PD- 1 is markedly upregulated on the surface of dysfunctional virus-specific CD8+ T cells during chronic HIV and SIV infections (Trautmann et al., 2006; Velu et al., 2009), and blockade of PD-1 *in vivo* enhanced SIV-specific CD8+ T cells responses (Velu et al., 2009). Moreover, recent studies found that high percentages of follicular CD8+ T cells in chronic HIV and SIV infection express inhibitory molecule PD- 1 (Li et al., 2016; Petrovas et al., 2017). However, the degree to which follicular and extrafollicular SIV- specific CD8+ T cells in early infection express PD-1 has not yet been investigated.

To determine the percentage of SIV-specific CD8+ T cells expressing PD-1 in follicular and extrafollicular compartments, we stained lymph node tissue sections from SIV infected rhesus macaques with MHC-class I tetramers, antibodies directed against PD-1, and antibodies directed against CD20 to label B cell follicles. We found PD-1+ SIV-specific tetramer+ CD8+ T cells in both follicular and extrafollicular areas in early SIV infection at 21 dpi (Fig. 3A). Quantitative analysis showed a broad range of both follicular (median 62%; range 12-67%) and extrafollicular (median 65%; range 8-68%) SIV- specific tetramer+ CD8+ T cells expressing PD-1. No significant difference was observed between the percentage of PD-1+ SIV-specific tetramer+ CD8+ T cells inside and outside B cell follicles (Fig. 3B). We further compared the percentage of SIV-specific tetramer+ CD8+ T cells that express PD-1 between early and our previously published data (Li et al., 2016) from chronic SIV infection in follicular and extrafollicular regions respectively. Again, no significant differences were observed (Fig. 3C and 3D).

**FIG 3.**
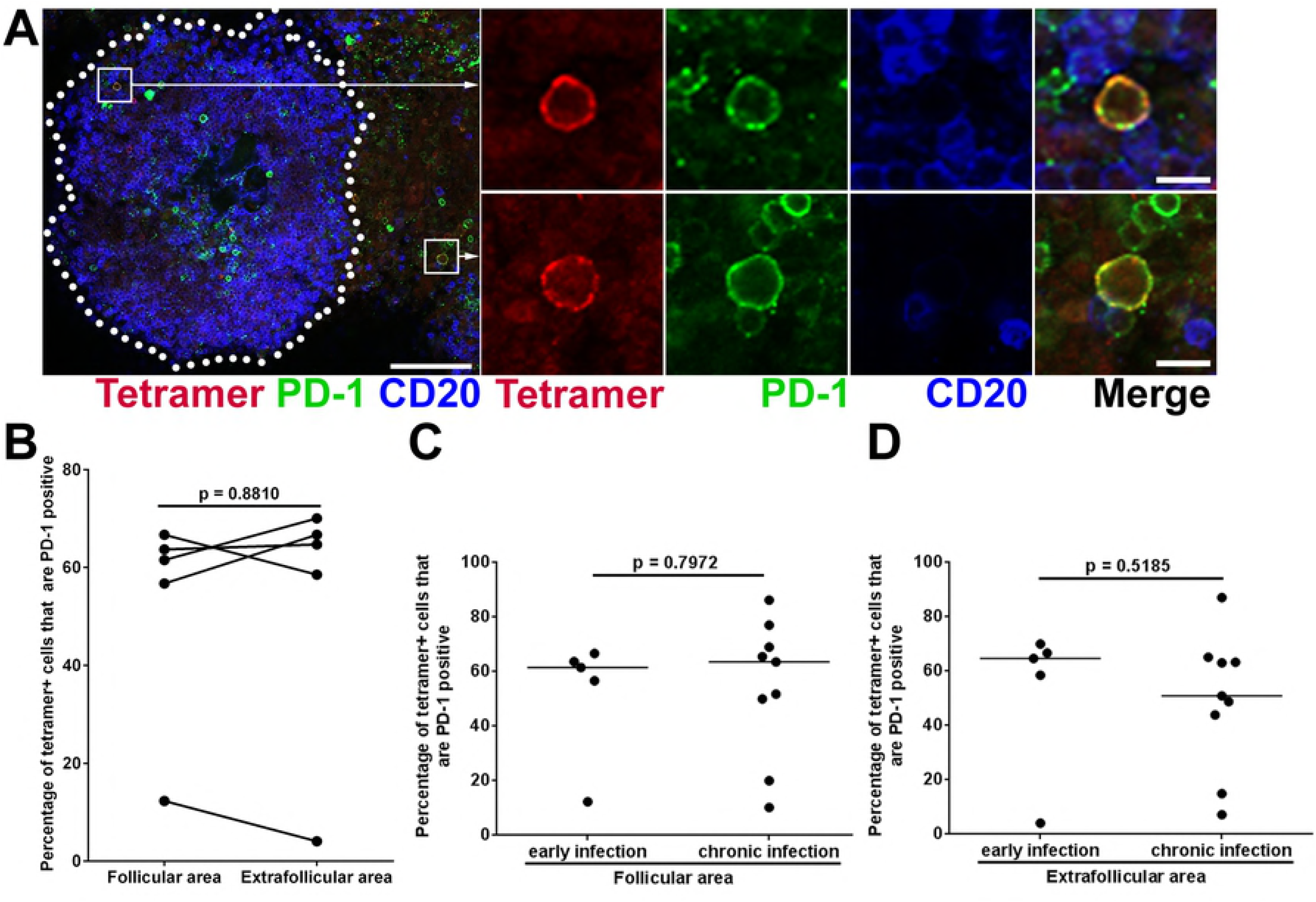
Subsets of tetramer+ SIV-specific CD8 T cells expressed PD-1 during early SIV infection. (**A**) Representative lymph node section shows tetramer+ SIV-specific CD8 T cells express PD-1 inside and outside B cell follicle. This section was stained with Mamu-A*001:01/Gag CM9 tetramers to label SIV-specific CD8+ T cells (red), PD-1 antibodies (green) to label PD-1 expressing cells and CD20 antibodies (blue) to define follicles. Confocal images were collected with a 20X objective and the scale bar is 100 µm in the image on the left and 10 µm in the enlargement. (**B**) There was no significant difference between the percentages of PD-1+ cells within the tetramer-binding population located in follicular and extrafollicular regions during early infection. There is no significant difference between percentages of tetramer+ SIV-specific CD8 T cells that are PD-1+ during early and chronic SIV infection in both (**C**) follicular (p = 0.7972) and (**D**) extrafollicular area (p = 0.5185).

### Foxp3+ T reg cells may inhibit follicular SIV-specific CD8+ T cells during early infection

Regulatory T cells (Tregs) play a crucial role in maintaining immunological self-tolerance and controlling autoimmune diseases (Sakaguchi et al., 2008). They also are involved in suppressing immune activation in viral infection (Sakaguchi et al., 2009; Schmidt et al., 2012). A large proportion of Tregs are characterized by the expression of the transcription factor Foxp3 (Hori et al., 2003; Fontenot et al., 2003). While most Tregs are CD4+, there exist a small population of CD8+ Tregs (Suzuki et al., 2012; Nigam et al., 2010). Directed contact is an important mechanism mediates suppression by Tregs (Park et al., 2015).

We next investigated whether Foxp3+ cells were in contact with and potentially inhibiting function of follicular SIV-specific CD8+ T cells in early infection. We stained lymph node tissue sections from rhesus macaques in early stage of SIV infection, at 21 dpi, with MHC-class I tetramers to label SIV- specific CD8+ T cells, anti-Foxp3 antibodies to label Foxp3+ Tregs, and anti-IgM antibodies to label B cell follicles. We found follicular SIV-specific tetramer+ CD8+ T cells that directly contact Foxp3+ cells (Fig. 4A) and follicular SIV-specific tetramer+ CD8+ T cells that expressed Foxp3+ (Fig. 4B). An average of 12.4% (range 7-20%) follicular SIV-specific tetramer+ CD8+ T cells were in direct contact with Foxp3+ cells and the corresponding number in extrafollicular was 18.6% (rang 9-26%) (Fig. 4C). No significant difference was shown between the percentage of SIV-specific tetramer+ CD8+ T cells that contact Foxp3+ cells inside and outside B cell follicles in early SIV infection (Fig. 4C). The percentages of Foxp3+ SIV-specific tetramer+ CD8+ T cells inside and outside B cell follicles also showed no significant difference (Fig. 4D).

**FIG 4.**
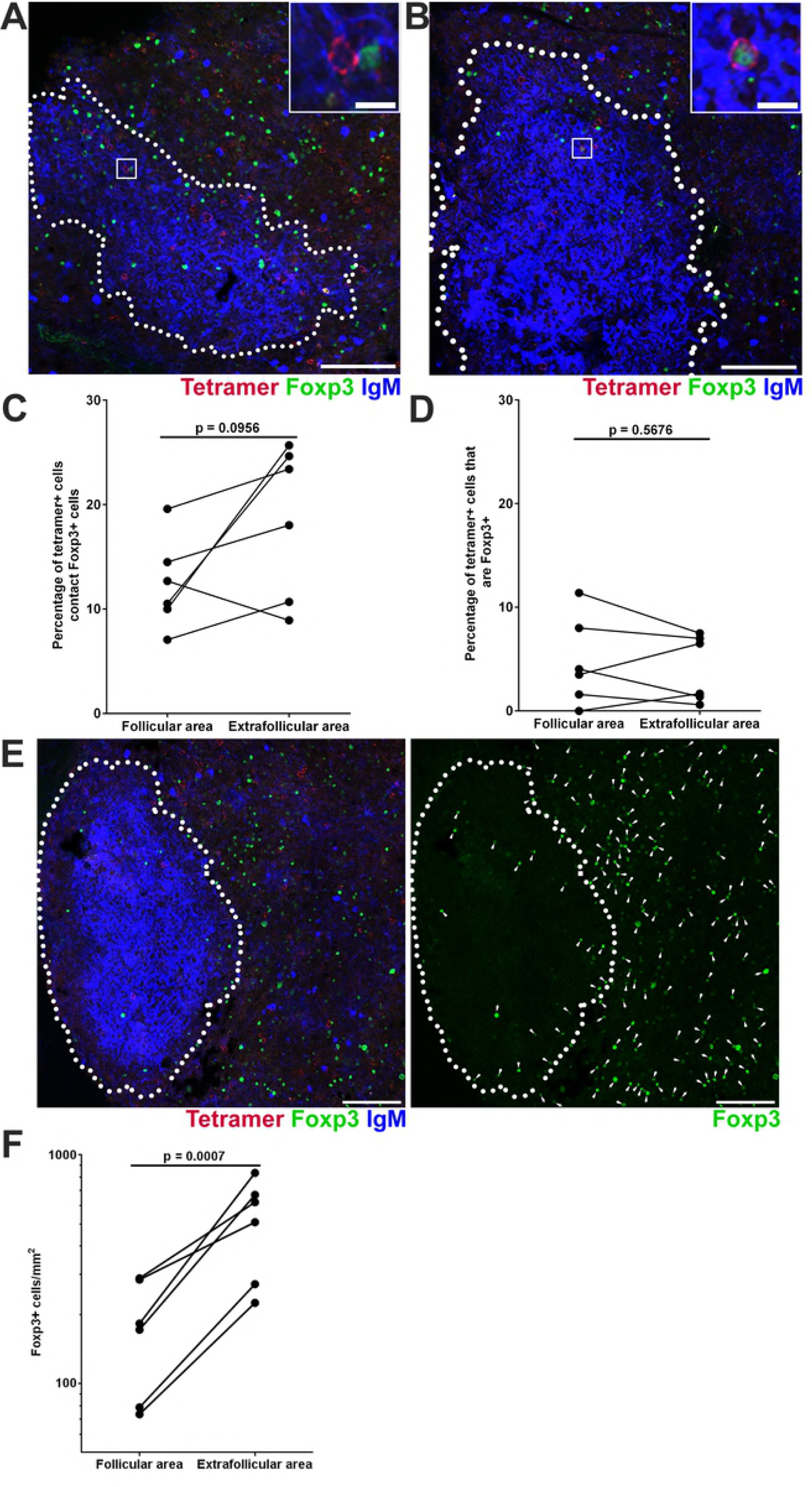
Follicular and extrafollicular tetramer+ SIV-specific CD8 T cells in relation to Foxp3+ cells during early SIV infection. (**A**) Representative lymph node section stained with Mamu-A*001:01/Gag CM9 tetramers (red), IgM (blue), and Foxp3 (green) showing tetramer+ cells in contact with Foxp3+ cells. Confocal images were collected with a 20X objective and the scale bar is 100 µm and 10 µm in low- and high-magnification images respectively. (**B**) Representative image showing tetramer+ cell is Foxp3+. (**C**) Percentages of follicular tetramer+ SIV-specific CD8 T cells that were in direct contact with Foxp3+ cells tend to be lower than extrafollicular tetramer+ SIV-specific CD8 T cells (95% CI: -13.9, 1.57; p = 0.0956). **(D**) There was no significant difference between percentages of tetramer+ SIV-specific CD8+ T cells inside and outside follicle that were Foxp3+ (95% CI: -2.1, 3.4; p = 0.5676). (**E**) Representative image showing distribution of Foxp3+ cells in lymph node. Scale bar is 100 µm. (**F**) Frequencies of Foxp3+ cells inside follicles were 67% (95% CI: 51%, 77%) lower than outside of follicle (p = 0.0007).

Similar to SIV-specific CD8+ T cells, Foxp3+ cells levels were also significantly lower in follicular than extrafollicular regions (Fig. 4E and 4F). In addition, there was no significant difference between the ratios of Tet+ cells: Foxp3+ cells in follicular and extrafollicular regions (data not shown). These findings suggest that contact mediated suppression of Foxp3+ Tregs on SIV-specific CD8+ T cells is similar in follicular and extrafollicular regions in early SIV infection.

We next evaluated whether the relationship of Foxp3+ Tregs and SIV-specific CD8+ T cells changed from early stages of infection to chronic infection. Interestingly, we first found significantly higher levels of Foxp3+ cells in early infection compared to chronic infection (Li et al., 2016) in follicular areas (Fig. 5A), but not in extrafollicular areas (Fig. 5B). Second, the percentage of follicular SIV-specific tetramer+ CD8+ T cells in contact with Foxp3+ cells in early infection was significantly higher than chronic infection (Fig. 5C), and this difference was not observed in extrafollicular SIV-specific CD8+ T cells (Fig. 5D). Third, the percentage of SIV-specific tetramer+ CD8+ T cells in follicular areas that expressed Foxp3 was significantly higher during early infection than chronic infection in both follicular (Fig. 5E) and extrafollicular areas (Fig. 5F). Lastly, the ratios of Tet+ cells: Foxp3+ cells in both follicular and extrafollicular regions were significantly lower in early infection compared to chronic infection (Fig. 5G and 5H).

**FIG 5.**
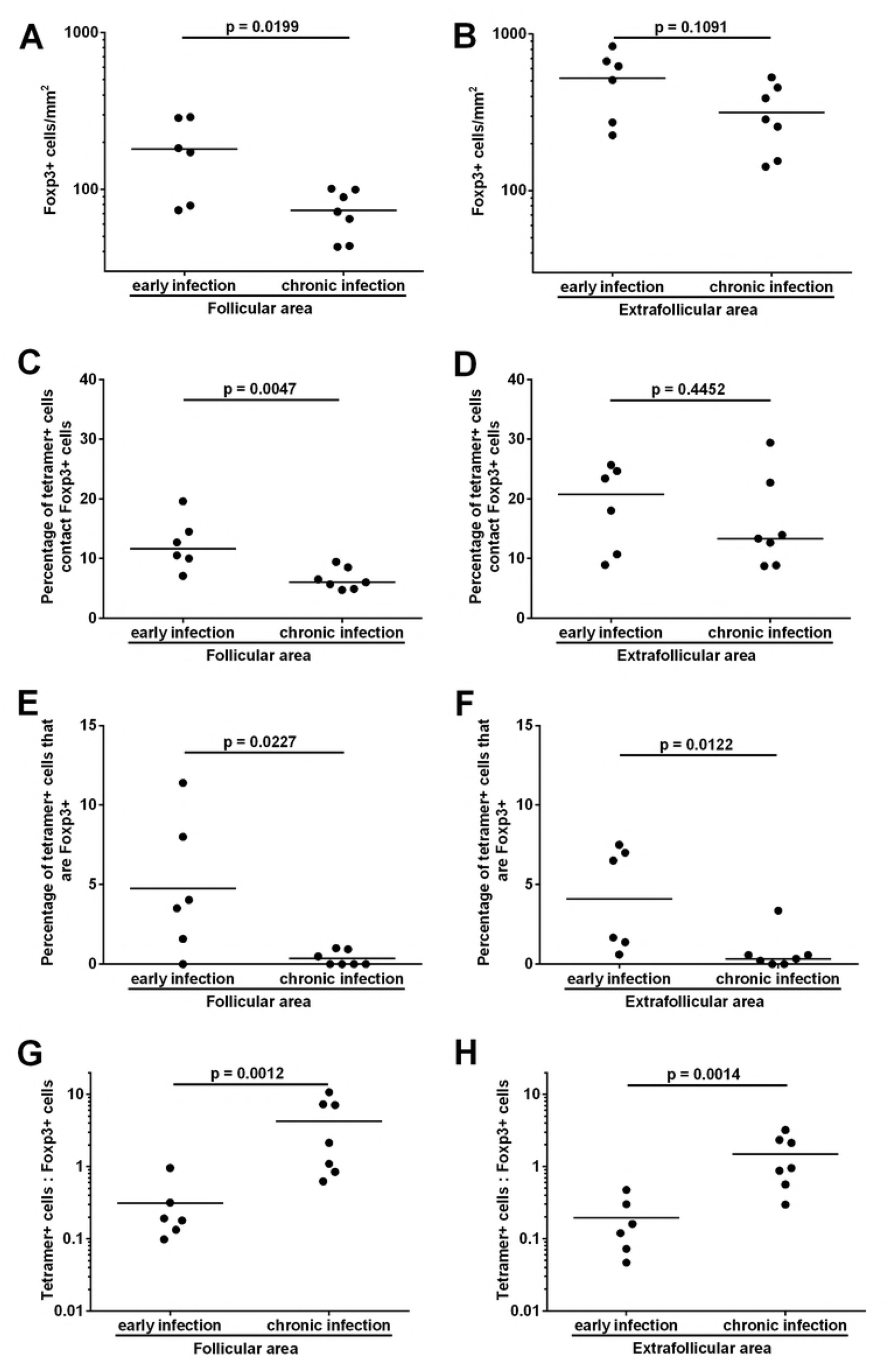
Foxp3+ Tregs relationship to follicular and extrafollicular SIV-specific CD8 T cells in early verses chronic SIV infection. There was significantly higher frequency of Foxp3+ cells during early SIV infection than chronic SIV infection in (**A**) follicular areas (2.26 times higher, 95% CI: 1.18, 4.34; p = 0.0199), but not in (**B**) extrafollicular areas (1.66 times higher, 95% CI: 0.88, 3.14; p = 0.1091). Significantly higher percentage of tetramer+ SIV-specific CD8 T cells during early SIV infection contact Foxp3+ cells than those during chronic SIV infection in (**C**) follicular areas (p = 0.00047), but not in (**D**) extrafollicular areas (p = 0.4452). (**E**) Percentage of follicular tetramer+ SIV-specific CD8 T cells that are Foxp3+ cells during early SIV infection are significantly higher than chronic infection (p = 0.0227). (**F**) At the same time, the percentage of extrafollicular tetramer+ SIV-specific CD8 T cells that are Foxp3+ cells during early SIV infection is also significantly higher than chronic infection (p = 0.0122). Ratios of tetramer+ SIV-specific CD8 T cells to Foxp3+ cells during early SIV infection were significantly lower than chronic SIV infection in both (**G**) follicular (91% lower; 95% CI: 70%, 97%; p = 0.0012) and (**H**) extrafollicular areas (87% lower; 95% CI: 63%, 96%; p = 0.0014).

### Activated proliferating follicular SIV-specific CD8+ T cells detected during early infection

We next assessed the levels SIV-specific CD8+ T cells that were activated and proliferating in follicular and extrafollicular regions of lymph nodes during early SIV infection, at 21 dpi. Ki67 is an activation and proliferation marker of T cells (Scholzen and Gerdes, 2000; Soares et al., 2010). Ki67+ SIV-specific tetramer+ CD8+ T cells were found in follicular as well as in extrafollicular regions (Fig.6A). Quantitative image analysis showed that the percentage of follicular Ki67+ SIV-specific tetramer+ CD8+ T cells was significantly lower than in extrafollicular counterparts (Fig. 6B). An average of 40% (range 7-61%) of SIV-specific tetramer+ CD8+ T cells were Ki67+ in follicular area compared to 54% (range 19-76%) in extrafollicular areas. To evaluate whether there was a change in Ki67 expressing virus-specific CD8+ T cells during early compared to chronic infection, we compared these findings to our previously published data from the chronic stage of infection (Li et al., 2016). We found that significantly fewer SIV-specific tetramer+ CD8+ T cells in both follicular and extrafollicular regions during chronic SIV infection expressed Ki67 (Fig. 6C and 6D).

**FIG 6.**
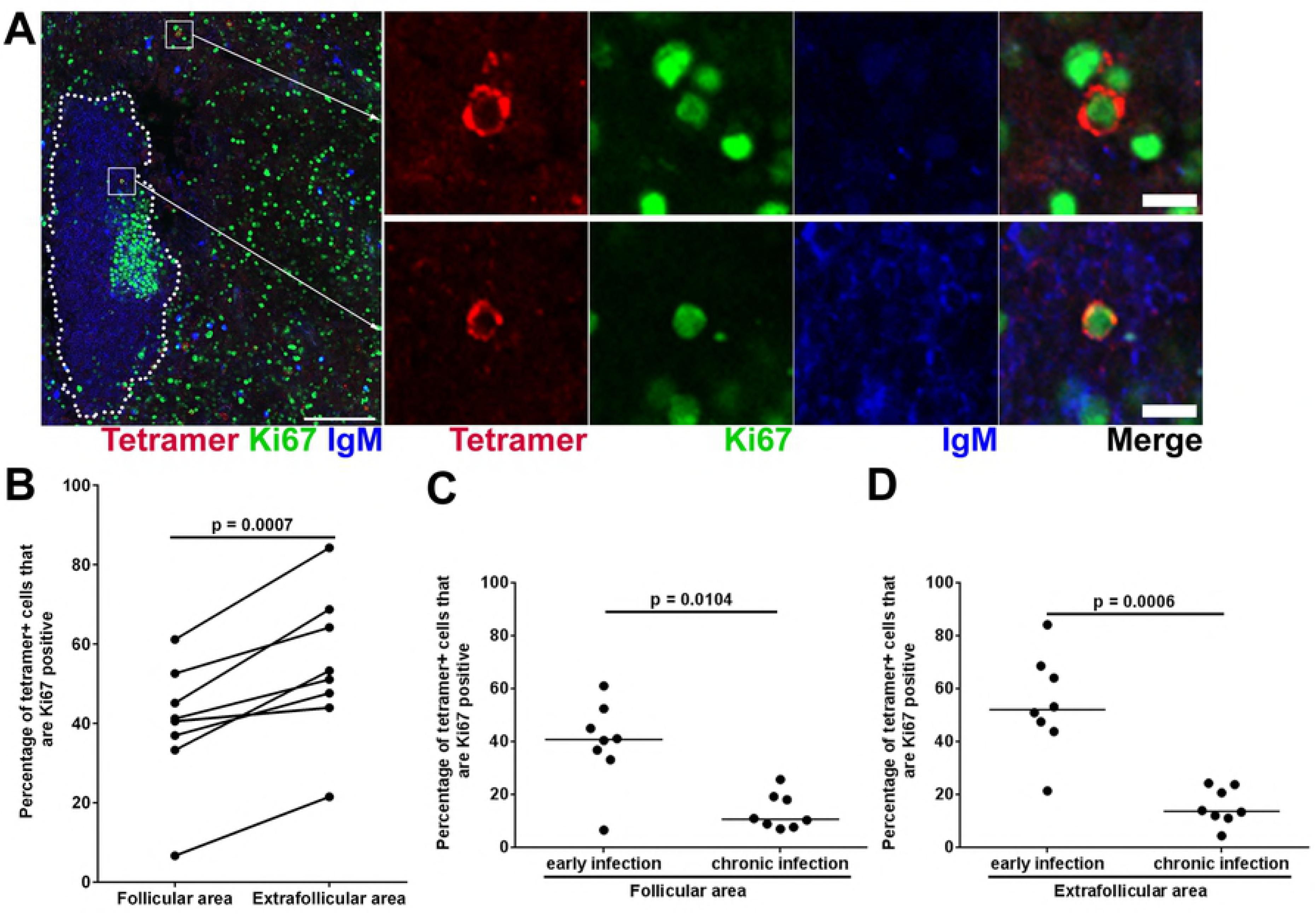
Ki67 expression levels in follicular and extrafollicular tetramer+ SIV-specific CD8 T cells. (**A**)Representative lymph node section stained with Mamu-A*001:01/Gag CM9 tetramers (red), IgM (blue), and Ki67 (green) showing tetramer+ Ki67+ cells in follicular and extrafollicular region. Scale bars indicate 100 µm and 10 µm in low- and high-magnification images respectively. (**B**) Percentages of tetramer+ SIV-specific CD8 T cells that expressed Ki67 in follicular areas is 14.6 percentage points (95% CI: 8.7, 20.6) lower than extrafollicular areas in lymph nodes during early SIV infection (p = 0.0007). Percentages of tetramer+ SIV-specific CD8 T cells that are Ki67+ during early SIV infection were significantly higher than chronic SIV infection in both (**C**) follicular (p = 0.0104) and (**D**) extrafollicular areas (p = 0.0006).

### Most follicular SIV-specific CD8+ T cells express perforin during early infection

Perforin is a crucial factor for cytolytic function in virus-specific CD8+ T cells. We previously showed that during chronic infection approximately 35% of follicular SIV-specific CD8+ T cells express perforin and that most expressed another cytolytic effector molecule, granzyme B, which typically works in concert with perforin to lyse infected cells (Connick et al., 2014; Li et al., 2016). Here we determined the expression of perforin in SIV-specific CD8+ T cells during early infection. Perforin expression levels within SIV-specific CD8+ T cells were divided into four categories (negative, low, medium, and high) (Fig. 7A). We found that most follicular SIV-specific tetramer+ CD8+ T cells expressed perforin (mean 73%; range 35-94%). Similar levels of low, medium and high perforin expressing SIV-specific tetramer+ CD8+ T cells were observed in follicular and extrafollicular areas (Fig. 7B). Comparison of perforin expression in SIV-specific tetramer+ CD8+ T cells in early and chronic infection (Li et al., 2016), showed significantly higher levels of perforin+ SIV-specific tetramer+ CD8+ T cells in early SIV infection in both follicular (Fig. 7C) and extrafollicular regions (Fig. 7D). In particular, the percentage of perforin^high^SIV-specific tetramer+ CD8+ T cells were higher during early compared to chronic stages of infection (Fig. 7E and F).

**FIG 7.**
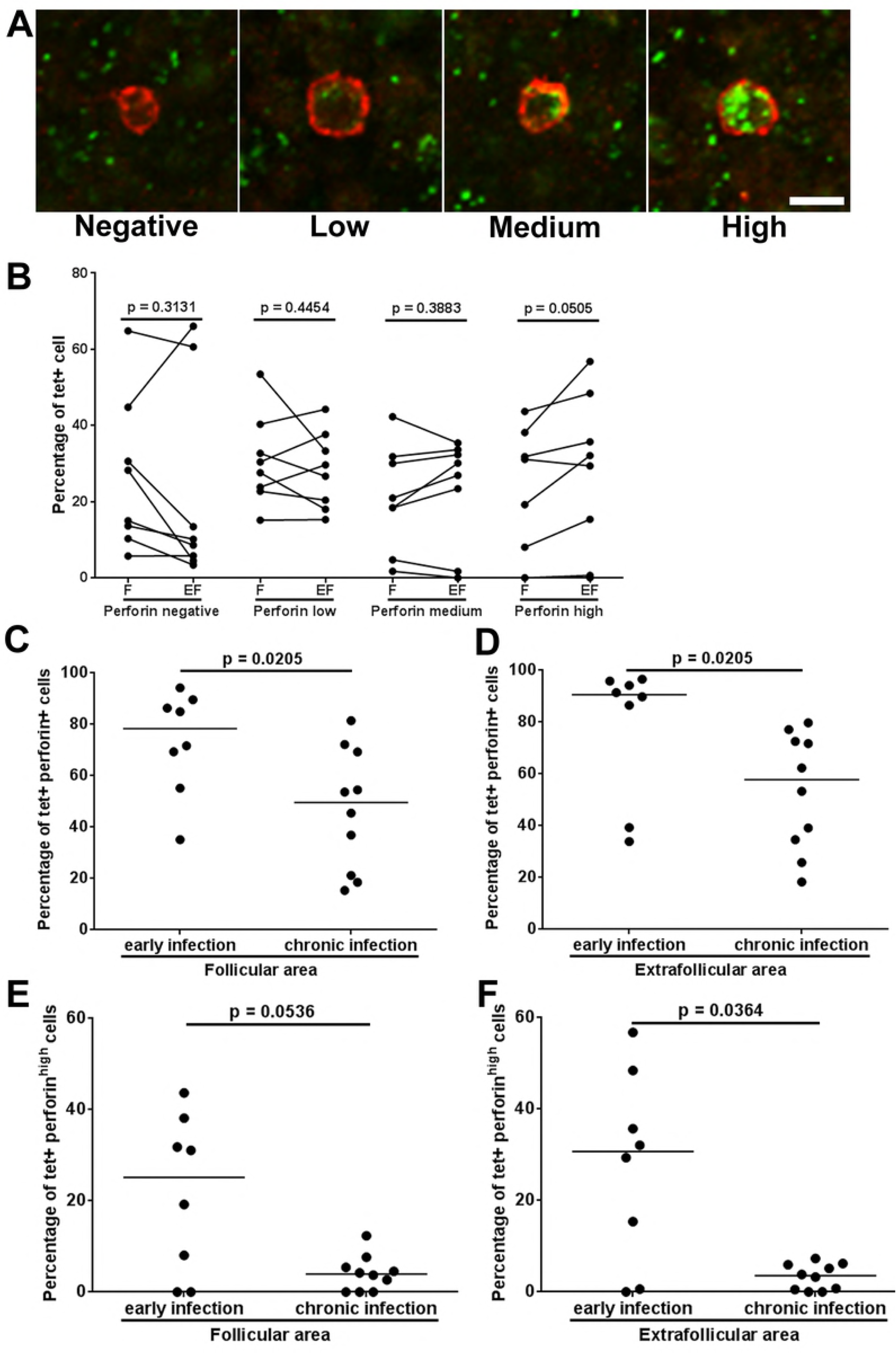
Most tetramer+ SIV-specific CD8 T cells expressed perforin during early SIV infection. (**A**) Representative lymph node section stained with Mamu-A*001:01/Gag CM9 tetramers (red) and perforin (green) showing perforin negative, perforin low, perforin medium and perforin high MHC-class I tetramer+ SIV-specific CD8 T cells. Scale bar indicates 10 µm. (**B**) Percentages of tetramer+ SIV-specific CD8 T cells that expressed perforin in follicular and extrafollicular regions. Among tetramer+ SIV- specific CD8 T cells, the distribution of cells across perfroin negative, low, medium and high is not significantly different between follicular and extrafollicular regions. Significantly higher percentage of tetramer+ SIV-specific CD8 T cells during early SIV infection express perforin than those during chronic SIV infection in both follicular area (p = 0.0205) (**C**), and in extrafollicular area (p = 0.0205) (**D**). Simultaneously, the percentages of tetramer+ SIV-specific CD8 T cells during early SIV infection tend to express a higher level of perforin than those during chronic SIV infection in follicular regions (p = 0.0536) (**E**), and are statistically significant in extrafollicular regions (p = 0.0364) (**F**).

### Cell death is not likely mediating low levels of follicular SIV-specific CD8+ T cells

A hypothesis yet to be tested regarding the relative low abundance of virus-specific CD8+ T cells in follicles is that follicular virus-specific CD8+ T cells die via apoptosis at greater rates than extrafollicular virus-specific CD8+ T cells. Here we tested that hypothesis and determined the levels of SIV-specific CD8+ T cells in follicular and extrafollicular levels that expressed Poly (ADP-ribose) polymerase (PARP) in early chronic SIV infection (50-60 dpi). PARP is involved in cells death by promoting release of apoptosis-inducing factor (AIF), and can be used as a marker of cells undergoing apoptosis (Yu et al., 2006). We found small subsets of SIV-specific tetramer+ CD8+ T cells that expressed PARP in both follicular as well as extrafollicular areas (Fig. 8A). Quantitative analysis showed no significant difference between the levels of PARP+ SIV-specific tetramer+ CD8+ T cells inside and outside B cell follicles (Fig. 8B).

**FIG 8.**
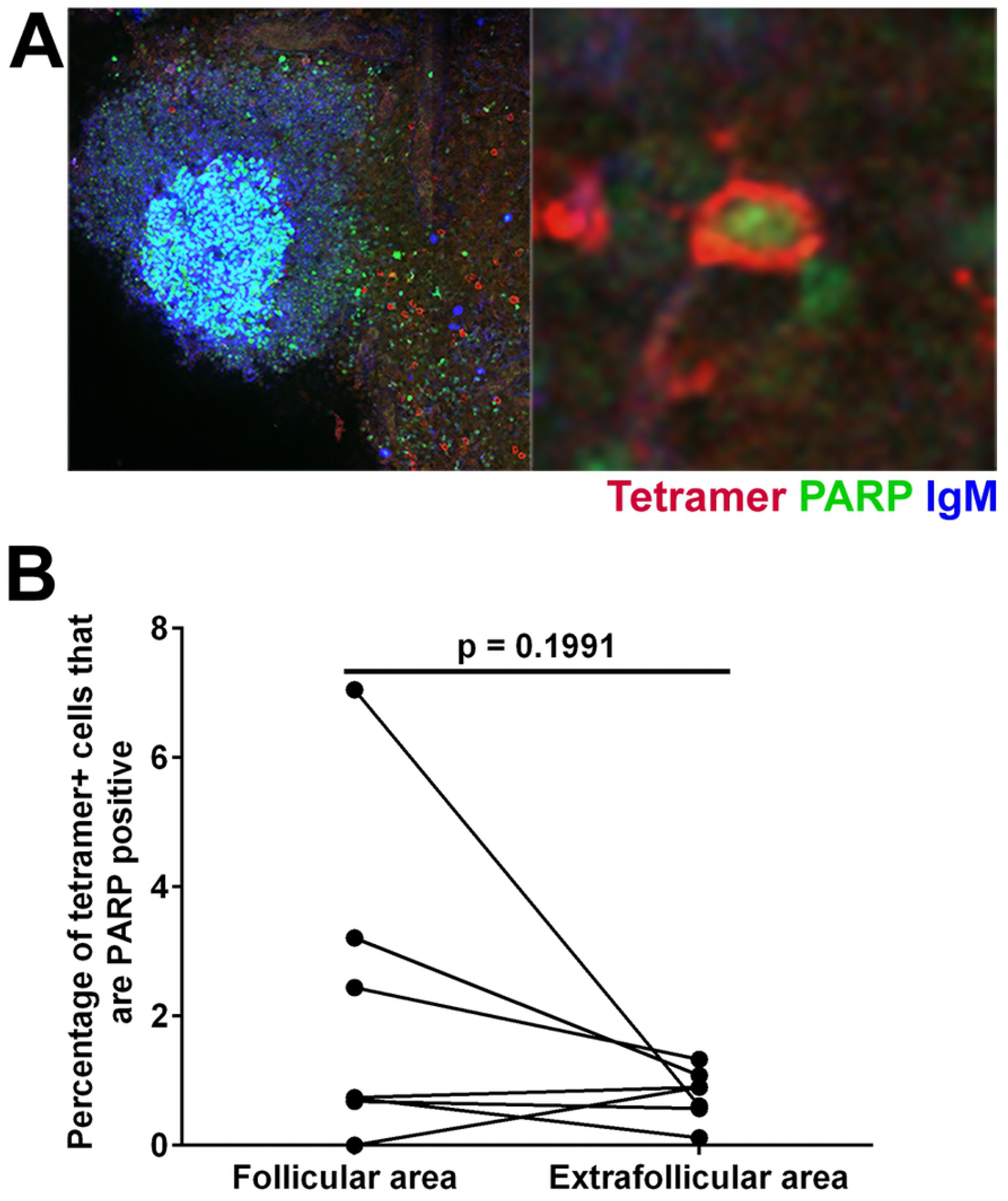
Subsets of of tetramer+ SIV-specific CD8 T cells in early chronic SIV infection express the apoptosis marker PARP. (**A**) Representative lymph node section stained with Mamu-A*001:01/Gag CM9 tetramers (red), IgM (blue), and PARP (green) showing tetramer+ PARP+ cells and tetramer+ cells in contact with PARP+ cells. Confocal images were collected with a 20X objective and the scale bar is 100 µm and 10 µm in low- and high-magnification images respectively. (**B**) The percentage of tetramer+ SIV-specific CD8 T cells that are PARP+ is not significantly different between follicular and extrafollicular regions (95% CI: -0.9, 3.6; p = 0.1991).

### Follicular SIV-specific CD8+ T cells likely suppress follicular SIV during early infection

In order to gain evidence as to whether levels of SIV-specific CD8+ T cells were impacting viral replication in follicular and extrafollicular compartments, we compared levels of SIV-specific tetramer+ CD8+ T cells and SIV RNA+ cells in follicular and extrafollicular regions of lymph nodes. In situ hybridization was used to localize and quantify SIV RNA+ cells and in situ tetramer staining used to localize and quantify cells at 21 dpi. We found that SIV-specific tetramer+ CD8+ T cells were significantly inversely correlated with SIV RNA+ cells in follicular regions (Fig. 9A), but not in extrafollicular regions (Fig. 9B).

**FIG 9.**
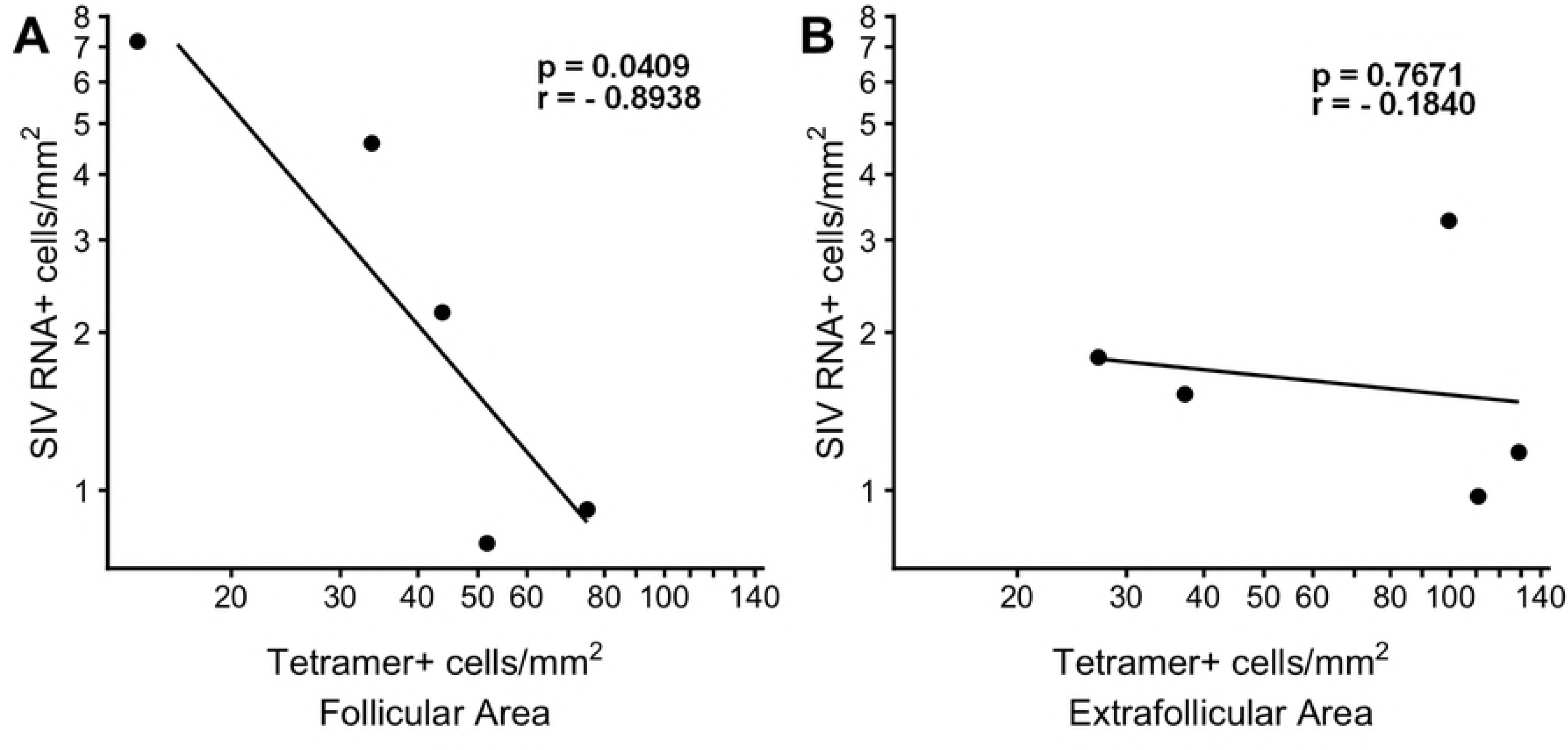
*In vivo* relationships between tetramer+ SIV-specific CD8 T cells and SIV RNA+ cells within F and EF compartments of lymph node at 21dpi. Frequencies of tetramer+ SIV-specific CD8 T cells significantly inversely correlated with SIV RNA+ cells in follicular region (p = 0.0409); a 1-log increase in tetramer+ SIV-specific CD8 T cells is associated with a 1.38-log (95% CI: 0.11, 2.65) decrease in SIV (**A**); but not in extrafollicular region (p = 0.7671), a one-log increase is tetramer+ SIV-specific CD8 T cells is associated with a 0.12 log decrease is SIV (95% CI: -1.3, 1.1) (**B**).

## Discussion

Eradication of HIV-infected cells *in vivo* remains a critical obstacle to cure HIV infection. B cell follicles are major anatomical reservoirs that permit active HIV and SIV replication during chronic infection (Connick et al., 2007, 2014; Fukazawa et al., 2015). However, virus-specific CD8+ T cells generally fail to accumulate in high frequency in B cell follicles (Connick et al., 2007, 2014; Li et al., 2016; Tjernlund et al., 2010). The ongoing viral replication in B cell follicles during chronic infection is likely due, at least partially, to the paucity of follicular anti-viral CD8+ T cell responses (Folkvord et al., 2005; Connick et al., 2007, 2014; Fukazawa et al., 2015; Skinner and Connick, 2014). It is not known whether this phenomenon also occurs during early stages of infection. Here, we determined the location, abundance, and phenotype of SIV-specific CD8+ T cells in follicular and extrafollicular regions during early SIV infection. We found few follicles had GCs at 14 dpi and abundant follicles with GCs at 21 dpi, suggesting that SIV infection had induced the formation of GCs as part of the immune response to combat SIV infection. At 21 dpi, we found that SIV-specific CD8+ T cells were largely excluded from B cell follicles, similar to what we reported during chronic disease (Connick et al., 2014). Strikingly, and in contrast to chronic infection, we also discovered that follicular SIV-specific CD8+ T cells were entirely absent in 60% of GC areas examined. A complete absence of SIV-specific CD8+ T cells in most GCs at 21 dpi whilst during chronic infection there are equal numbers of SIV-specific CD8+ T cells in GC and non-GC follicular areas (Li et al., 2016) suggests that during early infection, there is a temporal delay from the formation of nascent follicles and the entry of SIV-specific CD8+ T cells. Importantly, by 21 days dpi, the time in which we found most GC areas examined to be entirely devoid of SIV-specific CD8+ T cells, SIV has already seeded FDCs in GCs (Miller et al., 2005). These findings indicate that the arrival of SIV-specific CD8+ T cells in GCs occurs after SIV has already seeded FDCs and infected follicular CD4 T cells, thus setting the stage for the establishment of persistent chronic infection

In addition to determining the quantity of SIV-specific CD8+ T cells in follicular and extrafollicular compartment of lymph nodes during early infection, we also determined the phenotype of these cells. We found a broad range of SIV-specific CD8+ T cells expressing PD-1 during early infection. The PD-1 expression on SIV-specific CD8+ T cells during early infection likely reflects cells that were recently were exposed to antigenic stimulation, however, exhaustion cannot be ruled out. We found subsets of SIV-specific CD8+ T cells were in direct contact with FoxP3+ cells and some SIV-specific CD8+ T cells were FoxP3+. These data suggest that the function of some follicular as well as extrafollicular SIV-specific CD8+ T cells may be inhibited by Foxp3+ cells during early SIV infection. However, since FoxP3 is transiently induced on T cells upon activation *in vitro* (Morgan et al., 2005), additional studies are needed to confirm inhibition. We found subsets of SIV-specific CD8+ T cells expressing the activation and proliferation marker Ki67 in both follicular as well as extrafollicular areas, with higher levels in extrafollicular areas. These data indicate that during early infection, virus-specific CD8+ T cells were activated and proliferating in both follicular as well as extrafollicular compartments; and that most activated and proliferating cells were located in the extrafollicular areas. Levels of SIV- specific CD8+ T cells expressing Ki67 were higher in early infection compared to those we found during chronic stages of infection, and may be reflective of the reduction of virus and antigen for the SIV- specific CD8+ T cells during the transition of early to chronic stages of infection.

Most SIV-specific CD8+ T cells expressed perforin in both follicular as well as extrafollicular areas of lymph nodes during early infection indicating that these subsets were likely were capable of immediately killing an SIV infected cell upon contact. We found similar levels of tetramer+ virus-specific CD8+ T cells expressing perforin low, medium, and high levels of perforin or not expressing perforin in follicular and extrafollicular areas. With perforin expression levels being a marker of effector and memory subsets, this suggests that levels of effector and memory subsets are maintained at similar levels in follicular and extrafollicular compartments. Decreases in tetramer+ SIV specific T cells expressing high levels of perforin from early to chronic infection likely reflected the transition from the effector dominant CD8+ T cell response transitioning into a primarily memory pool of cells.

In this study, we also found similar levels of SIV-specific CD8+ T cells in follicular and extrafollicular areas of lymph nodes expressing the apoptosis marker PARP. These findings suggest that the relative low abundance of virus-specific CD8+ T cells in follicular relative to extrafollicular regions of lymph nodes is not likely due to follicular virus-specific CD8+ T cells dying via apoptosis at increased rates compared to extrafollicular virus-specific CD8+ T cells.

Importantly, during early SIV infection, levels of follicular virus-specific CD8+ T cells inversely correlated with levels of follicular SIV replication (SIV RNA+ cells). These findings suggest that follicular virus-specific CD8+ T cells are not only important in controlling SIV replication during early infection, but also that number matter.

## Materials and Methods

### Tissues from animals in early SIV infection

Lymph nodes were obtained from captive-bred rhesus macaques of Indian origin infected with SIVmac239 intravenously (IV). Portions of fresh lymphoid tissues were immediately snap frozen in OCT and/or formalin fixed and embedded in paraf?n. Simultaneously, portions of fresh lymphoid tissues were also collected in RPMI 1640 medium with sodium heparin (18.7 U/ml) and shipped overnight to the University of Minnesota for *in situ* tetramer staining.

### *In situ* tetramer staining combined with Immunohistochemistry

*In situ* tetramer staining combined with immunohistochemistry was performed on fresh lymph tissue specimens shipped overnight, sectioned with a compresstome (Abdelaal et al., 2015) and stained essentially as previously described (Skinner et al., 2000; Li et al., 2017). Biotinylated MHC-class I monomers were loaded with peptides (National Institute of Health Tetramer Core Facility, Emory University, Atlanta GA) and converted to MHC-class I tetramers. Mamu-A1*001 molecules loaded with SIV Gag CM9 (CTPYDINQM) peptides (Allen et al., 1998) or irrelevant negative control peptides FV10 (FLPSDYFPSV) from the hepatitis B virus core protein. Fresh lymph node sections were incubated with MHC-class I tetramers (0.5 µg/ml) alone or along with goat-anti-human PD-1 Abs (1 µg/mL, polyclonal, R&D Systems). For secondary incubations, sections were incubated with rabbit-anti-FITC Abs (0.5µg/mL, BioDesign, Saco, ME) and mouse-anti-human Ki67 Abs (1:500 dilution, clone MM1, Vector), or mouse-anti-human perforin Abs (0.1 µg/mL, clone 5B10, Novacastra), or mouse-anti-human Foxp3 Abs (2.5 µg/mL, clone 206D, BioLegend) or mouse-anti-human CD20 Abs (0.19 µg/mL, clone L26, Novocastra), or mouse-anti-human PARP Abs (3.5 µg/mL, clone Asp214, Cell signaling). For the tertiary incubations, the sections stained with goat-anti-human PD-1 Abs were incubated with Cy3-conjugated donkey-anti-rabbit Abs (0.3 µg/mL, Jackson ImmunoResearch Laboratories, West Grove, PA), Alexa 488-conjugated donkey-anti-goat Abs (0.75 µg/mL, Jackson ImmunoResearch Laboratories), and Cy5- conjugated donkey-anti-mouse Abs (0.3 µg/mL, Jackson ImmunoResearch Laboratories). All other sections were incubated with Cy3-conjugated goat-anti-rabbit Abs (0.3 µg/mL, Jackson ImmunoResearch Laboratories), Alexa 488-conjugated goat-anti-mouse Abs (0.75 µg/mL, Molecular probes), and Dylight 649-conjugated goat anti-human IgM (0.3 µg/mL, Jackson ImmunoResearch Laboratories). Sections were imaged using a Zeiss LSM 800 confocal microscope. Montage images of multiple 512 × 512 pixels were created and used for analysis.

### Quantitative image analysis

For the determination of levels of SIV-specific CD8+ T cells and percentages of SIV-specific CD8+ T cells that co-expressed specific molecules, follicular areas were identi?ed morphologically as clusters of brightly stained closely aggregated IgM^+^ or CD20^+^ cells. Follicular and extrafollicular areas were delineated using ImageJ software. Areas that showed loosely aggregated B cells that were ambiguous as to whether the area was a follicle were not included. For PD-1 expression analysis, an average of 112 tetramer^+^ cells (range, 67-190) was analyzed in follicular regions and 213 (range, 117-272) in extrafollicular regions. For quantification of tetramer^+^ cells that were in contact with Foxp3^+^ cells and express Foxp3^+^, an average of 102 tetramer^+^ cells (range, 57-193) was analyzed in follicular regions and 298 (range, 168-560) in extrafollicular regions. For Ki67 expression analysis, an average of 133 tetramer^+^ cells (range, 30-246) was analyzed in follicular regions and 307 (range, 130-464) in extrafollicular regions. For perforin expression level analysis, an average of 97 tetramer^+^ cells (range, 22-193) was analyzed in follicular region and 276 (range, 98-530) in extrafollicular region. To determine levels of perforin expression, tetramer^+^ cells were scored using the following objective criteria as follows. Tetramer^+^ cells with no detectable perforin staining above background levels were scored as perforin negative. Tetramer^+^ cells with perforin staining 2-3X greater than background were scored as perforin low, with perforin staining 4-9X higher than background as perforin medium, and those with 10X or greater than background levels and with perforin staining detectable throughout much of the cytoplasm were scored as perforin high. Cell counts were done on single z-scans. While doing the cells counts, we demarcated cells using a software tool to avoid counting the same cell twice. All quantitative image analyses were done with lymph node tissues. An average of 7.42 mm^2^ (range, 5.63-10.08 mm^2^) was analyzed for each lymph node.

### *In situ* hybridization combined with immunohistochemistry

*In situ* hybridization for SIV RNA was performed as previously described (Connick et al., 2014). This technique identifies cells that are actively transcribing SIV, but not extracellular virions encapsulated in envelope glycoprotein and bound to FDC. Briefly, 6 µm frozen sections were fixed in 4% paraformaldehyde (Sigma-Aldrich, St. Louis, MO), hybridized overnight with digoxygenin labeled SIVmac239 antisense probes (Lofstrand Labs, Gaithersburg, MD) and visualized using NBT/5-bromo-4- chloro-3-indolyl phosphate (Roche, Nutley, NJ). Immunohistochemistry staining for B cells was performed in the same tissues using mouse-anti-human CD20 (clone 7D1; AbD Serotec, Raleigh, NC) and detected using HRP-labeled polymer anti-mouse IgG (ImmPressKit; Vector Laboratories, Burlingame, CA) and Vector NovaRed substrate (Vector Laboratories). SIV RNA^+^ cells were counted by visual inspection and classified as either inside or outside of B cell follicles which were identified morphologically as a cluster of CD20^+^ cells as previously described (Li et al., 2017). Total tissue area and area of follicles was determined by quantitative image analysis (Qwin Pro version 3.4.0; Leica, Cambridge, U.K.) and used to calculate the frequency of SIV+ cells per mm^2^. An average of 12.5 mm^2^ (7.1 mm^2^ – 87.2 mm^2^) was analyzed.

## Statistical analysis

To compare differences in cells/mm^2^, values were log-transformed and either paired t-tests or independent t-tests with unequal variance were performed, as appropriate; for reporting, estimates and 95% confidence intervals were back-transformed and reported as percent differences. For relationships between two measures of cells/mm^2^, values were also log-transformed and Pearson’s correlation and linear regressions were performed. To compare differences in percentages, paired t-tests or Wilcoxon rank-sum tests were performed, as appropriate. All tests were two-sided and p<0.05 was considered statistically significant. All calculations performed in R 3.4.3 (R Core Team, 2017).

### Ethics Statement

All animals were housed and cared for according to American Association for Accreditation of Laboratory Animal Care standards in accredited facilities. All animal procedures were performed according to protocols approved by the Institutional Animal Care and Use Committees of the Wisconsin National Primate Research Center.

The trained employees of the Animal Care Unit of the Wisconsin Primate Research Center provided daily care for the animals included in these studies. The Wisconsin Primate Research Center is AAALAC certified. Animals are cared for by a staff of veterinarians under the direction of Saverio Capuano, DVM who is the Attending Veterinarian, four staff veterinarians and trained vet techs. At least twice daily animals were evaluated for signs of pain, illness and stress observing appetite, stool, typical behavior etc. by the staff of Animal Care Unit. If any of the above parameters are found unacceptable a WPRC veterinarian will be contacted and appropriate treatments will be started. After a procedure, the affected site was observed for potential complications 24 hours later, or as directed by the veterinarian. An animal was euthanized at the recommendation of the attending WPRC veterinarian if any of the preceding conditions are observed and the veterinarians deem euthanasia necessary. Any deteriorating condition deemed particularly distressful to the animal (as assessed by research and/or veterinary staff) will be a condition for euthanasia.

Animals were treated for pain and discomfort for the effects of SIV infection on the advice of the veterinarian. Blood draws, leukapheresis, experimental SIV infections, and biopsies were done under anesthesia. Animals were anesthetized using ketamine (up to 15 mg/kg i.m.) to be reversed at the conclusion of a procedure by up to 0.25 mg/kg atipamizole (i.v. or i.m.). Monitoring of anesthesia recovery will be documented every 15 minutes until the animal is sitting upright, then every 30 minutes until the animal is fully recovered from the anesthesia.

Animals were monitored for disease progression. Disease progression is variable, thus we monitored and use multiple factors in the consideration of euthanasia. These factors include: weight loss of 20% of total body weight (to determine weight loss, animals were weighed at least every 90 days, according to Primate Center SOP 3.01, and more frequently if deemed necessary by research or veterinary staff), infection with an opportunistic pathogen and no response to treatment, progressive decline in condition regardless of treatment, chronic diarrhea and anorexia, neurological signs, such as disorientation, abnormal gait or posture, tremor etc. If any of the preceding conditions are observed and the veterinarians deem it necessary, the animal was euthanized. Any deteriorating condition deemed particularly distressful to the animal (as assessed by research and/or veterinary staff) was a condition for euthanasia. At the end of the study, or anytime during the study if recommended by a Center veterinarian, an animal will be euthanized. An animal will be euthanized by an IV overdose (greater than or equal to 50 mg/kg or to effect) of sodium pentobarbital or equivalent as approved by a clinical veterinarian, preceded by ketamine (at least 15 mg/kg body weight, IM). It is possible that the final blood draw, performed following anesthesia but prior to the IV overdose, may result in death by exsanguination. All euthanasia was performed in accordance with the recommendations of the Panel on Euthanasia of the American Veterinary Medical Association. Death would be defined by stoppage of the heart as determined by a qualified and experienced person using a stethoscope to monitor heart sounds from the chest area, as well as all other vital signs that can be monitored by observation. Necropsy will be performed by a qualified pathologist of the WNPRC.

## Acknowledgments

We thank the WNPRC pathologists and staff for care of animals and acquisition of tissues, the WNPRC Immunology Services for monitoring CD4 counts, and Virology Services for providing inoculum and monitoring plasma viral loads.

